# Synergistic Interaction Between Nitrofurantoin and Dequalinium Provides a Promising Approach to Treating UTIs caused by Antibiotic-Resistant *Klebsiella pneumoniae*

**DOI:** 10.64898/2026.08.03.742500

**Authors:** Eric Cheuk-Kiu Cheung, Catherine Stevens, Brynn Tate, Matthew A. Mulvey, Jessica C. S. Brown

## Abstract

Nitrofurantoin is commonly prescribed as a first-line treatment for urinary tract infections (UTIs) and is effective against most uropathogens; however, resistance among *Klebsiella* spp. remains high. In this study, we investigated synergistic drug combinations—defined as pairs of drugs whose combined effect exceeds that of each agent alone—to enhance treatment efficacy against drug-resistant *Klebsiella pneumoniae*. A high-throughput drug screen identified six and two small molecules that exhibited >50% synergistic activity in tested strains in combination with nitrofurantoin and ciprofloxacin, respectively. We further validated the top three candidates using nitrofurantoin-susceptible and resistant clinical isolates. Notably, the combination of nitrofurantoin and dequalinium demonstrated strong synergistic activity, observed in 68% of susceptible *Escherichia coli* strains and 94% of resistant *Klebsiella pneumoniae* strains tested. Combined inhibition of the citric acid cycle by nitrofurantoin and F1-ATPase by dequalinium resulted in a significant reduction in ATP levels compared to either treatment alone. Importantly, decreased ATP levels did not increase persister cell formation, and dequalinium alone reduced persister cell populations more effectively than nitrofurantoin. However, nitrofurantoin is processed into a poorly understood reactive intermediate whose specific metabolic targets have not been fully elucidated. Using metabolomic analyses, we identified aconitase and isocitrate dehydrogenase—key enzymes in the citric acid cycle—as altered in response to nitrofurantoin. Together, these findings demonstrate that dual targeting of bacterial metabolism by nitrofurantoin and dequalinium represents a promising therapeutic strategy for treating drug-resistant *Klebsiella* spp. in UTIs.

## Introduction

Urinary tract infections (UTIs) are among the most common infectious diseases worldwide, affecting over 400 million individuals annually and contributing to approximately 260,000 deaths each year [1]. UTIs are broadly classified as uncomplicated or complicated. Uncomplicated UTIs, which account for ∼60–65% of cases, typically occur as cystitis in otherwise healthy, immunocompetent, non-pregnant individuals without structural or functional abnormalities of the urinary tract [2]. In contrast, complicated UTIs (∼35–40%) are associated with factors such as pyelonephritis, urinary tract abnormalities, catheterization, or recurrent infections, and are often more difficult to treat [3–6].

UTIs are caused by a diverse range of pathogens, including Gram-negative and Gram-positive bacteria and fungi. However, the majority of infections are attributed to Gram-negative organisms, with *Escherichia coli* responsible for approximately 65–75% of cases. *Klebsiella spp.* represent the second most common causative agent and are of particular clinical concern due to their increasing prevalence and elevated rates of antibiotic resistance [5, 7, 8].

While uncomplicated UTIs are generally resolved with standard antibiotic therapy, complicated and recurrent infections often require repeated or prolonged treatment [6]. This repeated antibiotic exposure contributes to the emergence of resistance, leading to treatment failure and narrowing therapeutic options. Indeed, recurrent UTIs exhibit resistance rates of up to 80-90% to at least one antibiotic class and approximately 50-80% to three or more classes, underscoring the growing challenge of antimicrobial resistance in uropathogens [9, 10].

The development of new antibiotics has not kept pace with the rise in resistance. Antibiotic discovery is time-intensive, costly, and characterized by a high rate of candidate attrition [11, 12]. Notably, few truly novel antibiotic classes have been introduced in recent decades, with most new agents representing modifications of existing scaffolds [13, 14]. While such derivatives may improve potency or reduce toxicity, they often retain similar mechanisms of action, rendering them vulnerable to cross-resistance within the same class [14].

An alternative strategy to address antibiotic resistance is the identification of synergistic drug combinations [15, 16]. Synergy occurs when the combined effect of two compounds exceeds the sum of their individual effects, and in some cases can restore antibiotic activity against resistant strains [17, 18]. Clinically successful examples include trimethoprim-sulfamethoxazole (TMP–SMX). In the folic acid synthesis pathway, trimethoprim inhibits dihydrofolate reductase, which converts dihydrofolic acid to tetrahydrofolic acid, and sulfamethoxazole inhibits dihydropteroate synthase, which converts para-aminobenzoic acid to dihydrofolic acid. The inhibition of these sequential steps in the folic acid synthesis pathway allows the drugs to act synergistically [19–22]. These approaches extend the utility of existing antibiotics without requiring the development of entirely new drug classes.

In this study, we sought to identify synergistic combinations between antibiotics commonly used to treat UTIs and clinically approved small molecules. Drug repurposing offers a strategic advantage by leveraging compounds with established safety profiles, thereby reducing both development time and cost [23–25]. In addition to their therapeutic potential, synergistic interactions can provide mechanistic insight into antibiotic activity and bacterial physiology [24, 26, 27]. Here, we combine high-throughput screening with mechanistic analyses to identify and characterize synergistic drug pairs with activity against drug-resistant uropathogens.

## Results

### Nitrofurantoin and ciprofloxacin exhibit synergistic interactions with multiple small molecules in the US and International Drug Collection

To identify synergistic drug pairs, we selected two antibiotics commonly used to treat Gram-negative pathogens causing urinary tract infections (UTIs): nitrofurantoin (NFT), frequently prescribed for acute UTIs, and ciprofloxacin (CPFX), typically reserved for complicated cases [6, 28, 29]. NFT is used exclusively for UTIs due to its pharmacokinetic property of concentrating in the urine. While resistance to NFT remains relatively low (∼5%) in *Escherichia coli*, resistance in *Klebsiella pneumoniae* and *Klebsiella oxytoca* is substantially higher (∼40%) [30–32]. In contrast, CPFX is reserved for complicated UTIs due to high resistance rates (∼40–70%) [31–33]. These resistance profiles make NFT and CPFX suitable candidates for identifying synergistic partners.

To identify drugs that act synergistically with NFT or CPFX, we grew E. coli MG1655 in the presence of the Microsource Spectrum Drug Library, with or without target antibiotics NFT or CPFX. The Microsource Spectrum Drug Library contains >2,500 small molecules approved for clinical use by U.S. and international regulatory agencies (MicroSource Discovery Systems, Inc., Gaylordsville, CT, USA) [34, 35], enabling rapid repurposing for antibacterial applications. Each test well contained a sub-inhibitory concentration of the test antibiotic and Microsource Spectrum Collection small molecule; the Microsource Spectrum Collection alone served as a control. We tested two sub-inhibitory concentrations (2- and 4-fold below the minimum inhibitory concentration [MIC]) were tested for both NFT and CPFX to capture a range of potential interactions. We expected weakly synergistic interactors to inhibit growth at 2-fold below MIC, whereas strongly synergistic pairs would do so at 4-fold below MIC.

We then incubated plates for 24 h and used OD₆₀₀ as a stand-in for growth. Wells without growth identified potential synergistic combinations. However, intrinsic antibacterial activity of the library molecules could mask synergistic interaction(s) with NFT or CPFX, compounds that also inhibited growth in control wells were retained for additional testing. In total, we identified 21 and 15 small molecules with potential synergy in combination with NFT and CPFX, respectively (Fig. 2A and 1A).

**Figure 1.**
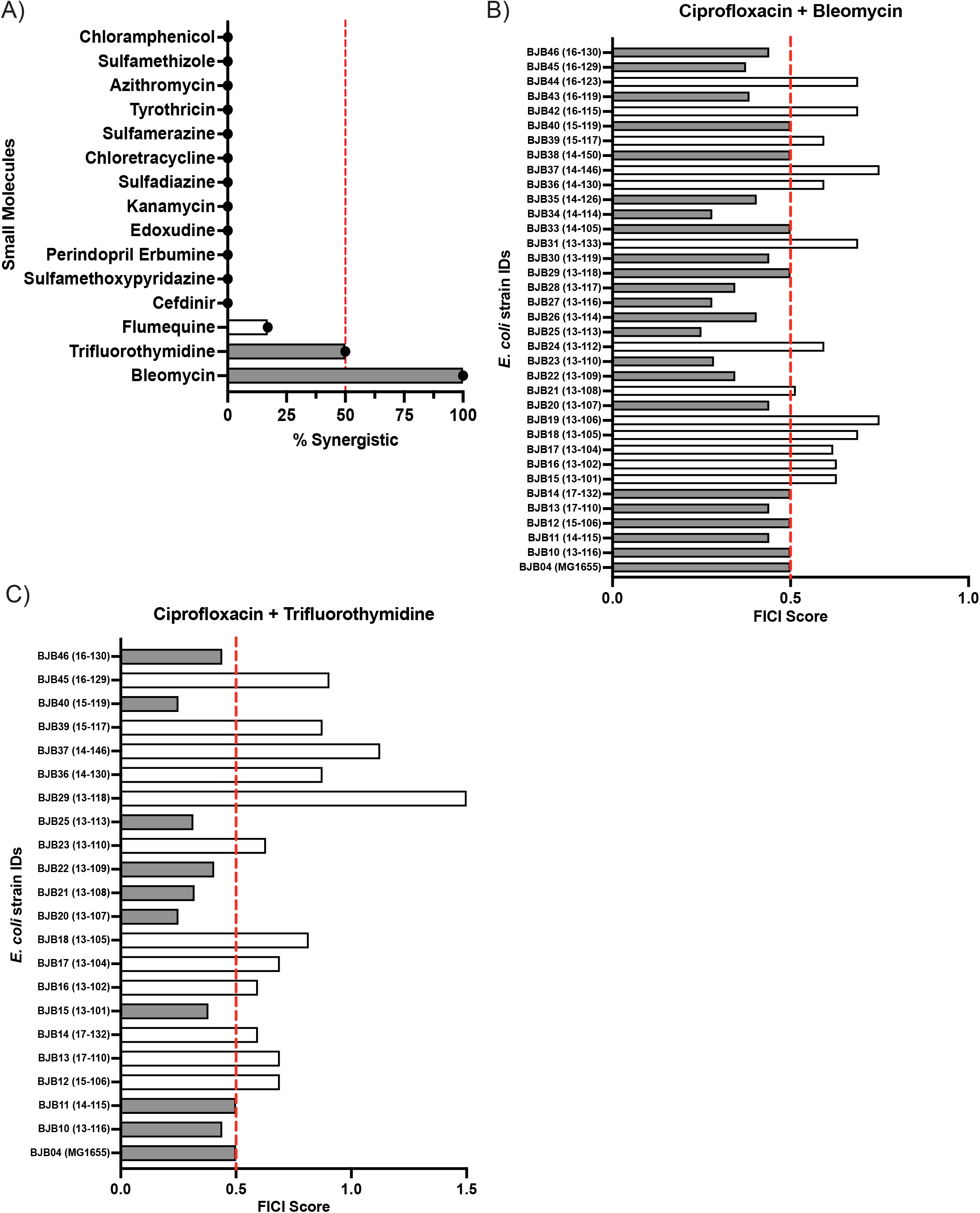
Evaluation of ciprofloxacin-based drug combinations identified from the high-throughput screen. **(A)** Initial synergy validation of small-molecule hits. Each drug combination was tested in triplicate using six susceptible *E. coli* strains, and the percentage of strains exhibiting synergy was graphed. Combinations were considered truly synergistic if they demonstrated synergy in >50% of the strains, indicated by bars to the right of the red dotted line. The top two combinations, represented by gray bars, were selected for further synergy testing using ciprofloxacin-susceptible *E. coli* clinical isolates. Synergy testing in these clinical isolates demonstrated synergy rates of **(B)** 64% for ciprofloxacin + bleomycin, and **(C)** 45% for ciprofloxacin + trifluorothymidine. All synergistic strains are represented by gray bars.

**Figure 2.**
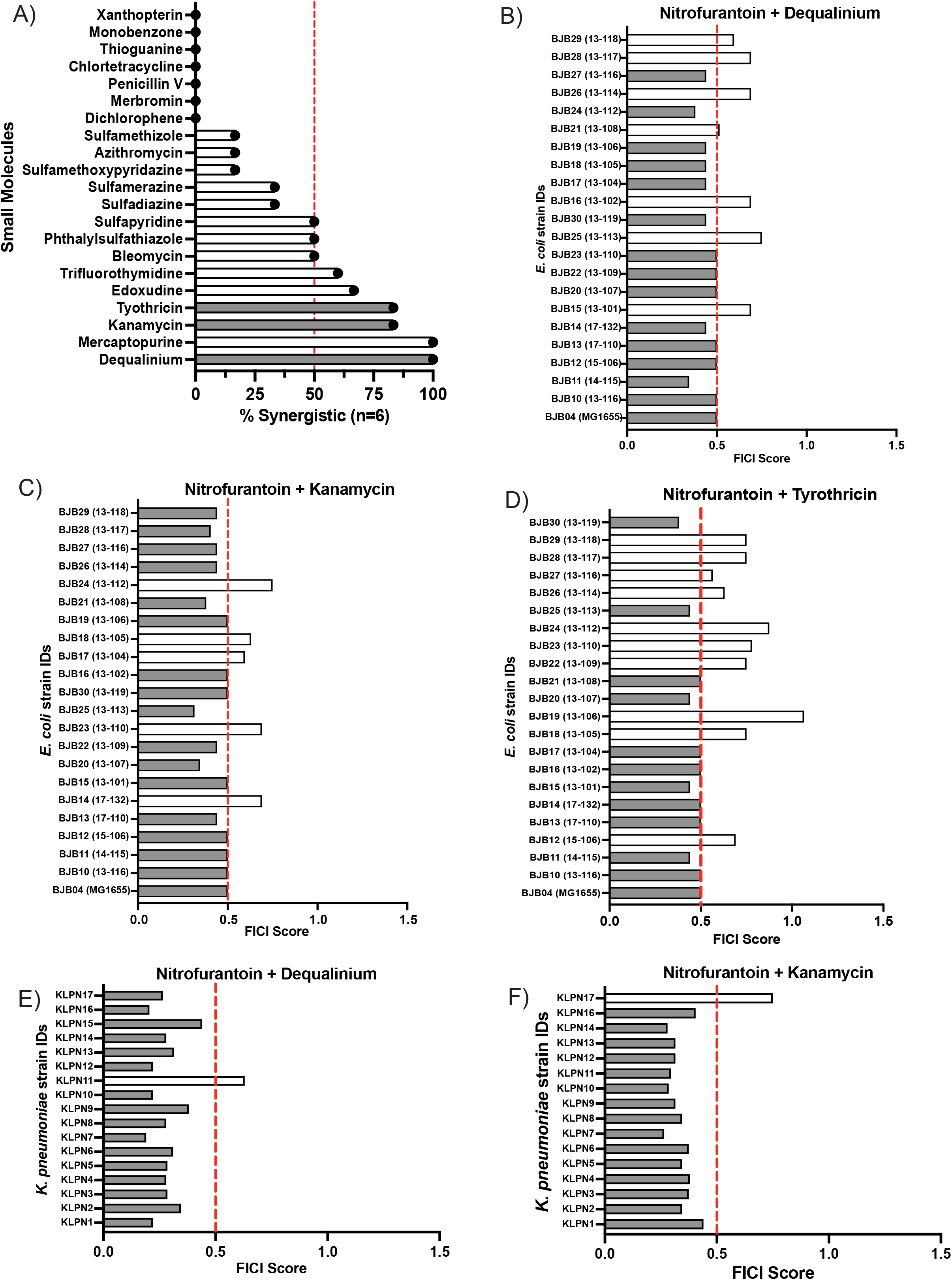
Evaluation of nitrofurantoin-based drug combinations identified from the high-throughput screen. **(A)** Initial synergy validation of small-molecule hits. Each drug combination was tested in triplicate using six NFT-susceptible *E. coli* strains, and the percentage of strains exhibiting synergy is shown. Combinations were considered synergistic if they demonstrated synergy in >50% of the strains, indicated by bars extending to the right of the red dashed line. The top three combinations (gray bars), excluding mercaptopurine due to cost, were selected for further synergy testing against NFT-susceptible *E. coli* clinical isolates. Checkerboard assay revealed synergy rates of **(B)** 68% for NFT + DEQ, **(C)** 77% for NFT + KAN, and **(D)** 55% for NFT + TYR. NFT + DEQ **(E)** and NFT + KAN **(F)** were also evaluated against NFT-resistant *K. pneumoniae* clinical isolates, with both combinations exhibiting synergistic activity in 94% of strains tested. Gray bars indicate synergistic strain pairs in all panels.

We validated all candidate synergistic combinations using the checkerboard assay, a standard method for assessing drug interactions. Briefly, 96-well plates were used to generate two-dimensional serial dilutions, with Drug A diluted across columns (1–11) and Drug B across rows (A–G). Column 12 and row H served as single-drug controls. After inoculation, plates were incubated at 37 °C for 24 h, and growth was quantified by OD₆₀₀. Wells exhibiting ≥90% inhibition were considered to show no growth. We assessed synergy by calculating the fractional inhibitory concentration index (FICI) 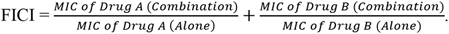 Combinations with FICI ≤ 0.5 were defined as synergistic, corresponding to at least a fourfold reduction in MIC for each drug in combination relative to monotherapy [23, 24, 36].

Our initial test for broad synergy involves testing *E. coli* MG1655 and four susceptible clinical *E. coli* isolates. For these first tests, we selected susceptible strains to minimize masking of weak synergistic effects. Combinations demonstrating synergy in ≥50% of tested strains were classified as synergistic and advanced for further analysis. Using this criterion, nine small molecules acted synergistically with NFT (Fig. 2) and two acted synergistically with CPFX (Fig. 1).

### Synergy is strongest in the nitrofurantoin–dequalinium and ciprofloxacin–bleomycin combinations

Following initial confirmation of synergistic interactions, we selected the top three NFT-based combinations and both CPFX-based combinations to further evaluate the extent of their activity. For NFT, dequalinium (DEQ), kanamycin (KAN), and tyrothricin (TYR) were prioritized for additional testing; mercaptopurine was excluded due to cost considerations. For CPFX, bleomycin (BLE) and trifluorothymidine (TRI) were selected, as they were the only two combinations that met the predefined criteria for synergy.

To assess clinical relevance, we tested these combinations against NFT- or CPFX-susceptible clinical isolates obtained from patients with positive urine cultures. Synergy was evaluated using the checkerboard assay. Among the NFT combinations, synergistic activity was observed in 68% (NFT+DEQ), 77% (NFT+KAN), and 55% (NFT+TYR) of the strains tested (Fig. 2B-D). For CPFX combinations, synergy was detected in 64% (CPFX+BLE) and 45% (CPFX+TRI) of strains (Fig. 1B-C).

Overall, our results demonstrate that the identified drug combinations retain synergistic activity across a diverse set of clinical isolates. However, for the remainder of this study, we focused on the synergistic drug pair nitrofurantoin-dequalinium. Kanamycin is already an FDA-approved antibiotic and therefore does not meet our aim of identifying a small-molecule that can be repurposed. We also excluded drug pairs containing ciprofloxacin, as its mechanism of action is well characterized. In contrast, NFT is a pro-drug that is converted into an active precursor and whose molecular target is poorly understood. We therefore sought to leverage synergistic drug pairs as a tool to better understand the mechanism of action of nitrofurantoin.

### The nitrofurantoin–dequalinium combination exhibits enhanced synergistic activity in nitrofurantoin-resistant isolates

We selected NFT because it is a first-line therapy for acute uncomplicated cystitis, which accounts for the majority of UTIs, and it exhibits broad antimicrobial activity against both Gram-positive and Gram-negative organisms [6, 29, 37, 38]. Treatment failure with NFT is often attributed to acquired resistance via the inactivation of nitroreductase enzymes or intrinsic resistance due to their absence [39–41]. Therefore, identifying a synergistic partner that restores nitrofurantoin efficacy against these resistant strains could significantly broaden its clinical utility. We selected DEQ due to its high synergistic activity. Furthermore, as it is not currently approved for use in the United States, demonstrating its effectiveness in this combination could support its repurposing for the domestic market [42].

Since our primary goal of this study was to identify combinations effective against resistant UTI pathogens, we next evaluated whether the observed synergy was maintained in NFT-resistant isolates. In the United States, resistance to nitrofurantoin in *E. coli* remains relatively low (∼5%), but higher rates (∼10%) have been reported globally [30–32, 43, 44]. Given this variability, we focused on *K. pneumoniae*, a common UTI pathogen responsible for approximately 10% of infections and characterized by substantially higher NFT resistance rates (∼40%) [5, 7, 10, 40].

Strikingly, we found that the NFT+DEQ combination exhibited enhanced synergistic activity in these resistant isolates, with synergistic response observed in 94% of strains tested (Fig. 2E). Only one isolate (KLPN11) did not show a synergistic response. We also tested the NFT+KAN combination and observed a synergistic response in 94% of strains tested (Fig. 2F). Although NFT+KAN showed slightly higher synergy than NFT+DEQ in our initial testing against susceptible strains (77% vs. 68%, respectively), it was reassuring to find that NFT+DEQ was equally effective against resistant isolates. Collectively, these findings indicate that the NFT+DEQ combination is highly effective against NFT-resistant *K. pneumoniae*., supporting its potential as a therapeutic strategy for treating resistant UTIs.

### The nitrofurantoin–dequalinium combination significantly reduces intracellular ATP levels

We then sought to understand the mechanism underlying this synergistic combination. Three primary hypotheses have been proposed to explain the mechanisms underlying drug synergy. First, the “gain-of-function” hypothesis suggests that the interaction between two agents produces a novel inhibitory activity not observed with either drug alone. Second, the “two-hit” or “parallel pathway” hypothesis posits that the drugs simultaneously inhibit distinct cellular processes, resulting in enhanced antimicrobial efficacy [26]. Third, the “bioavailability” hypothesis proposes that one compound increases the intracellular concentration or stability of the other, for example by enhancing uptake or reducing degradation [24, 27]. These potentially mechanisms are not mutually exclusive.

To investigate the mechanism of synergy between NFT and DEQ, we first reviewed the known properties of these compounds. Both agents are used as antimicrobials: NFT is approved in the United States, whereas DEQ is approved in several European countries but not in the United States. The mechanisms of action for both compounds are not fully defined and are likely multifactorial. NFT is reduced by bacterial nitroreductases (NfsA and NfsB) to generate reactive intermediates that damage multiple cellular components, including the cell wall, pyruvate metabolism, DNA via nonspecific binding, and ribosomal proteins [37]. However, the active intermediate generated from NFT is not well characterized [39, 45, 46]. DEQ has been reported to disrupt membrane integrity, induce protein denaturation, precipitate nucleic acids, and inhibit F1-ATPase [47, 48].

Given these diverse mechanisms, we hypothesized that the observed synergy arises from parallel pathway inhibition. Specifically, we propose that NFT disrupts enzymes within the tricarboxylic acid (TCA) cycle, while DEQ inhibits F1-ATPase, together resulting in impaired ATP production and depletion of cellular energy reserves.

To test this hypothesis, we quantified intracellular ATP levels using the Promega BacTiter-Glo assay. *E. coli* MG1655 and other clinical isolates were treated with sub-inhibitory concentrations of DMSO (control), NFT, DEQ, or the NFT+DEQ combination. All conditions were performed in triplicate, and mean values were analyzed. Both NFT and DEQ individually reduced ATP levels relative to the control, with no significant difference observed between the two treatments. In contrast, the NFT+DEQ combination resulted in a significantly greater reduction in ATP levels compared to either agent alone (Fig. 3), albeit not greater than the sum of the ATP level reductions of each single agent. Our results indicate that combined treatment with NFT+DEQ leads to pronounced depletion of intracellular ATP, supporting a model in which concurrent disruption of metabolic pathways and ATP synthesis underlies their synergistic antibacterial activity.

**Figure 3.**
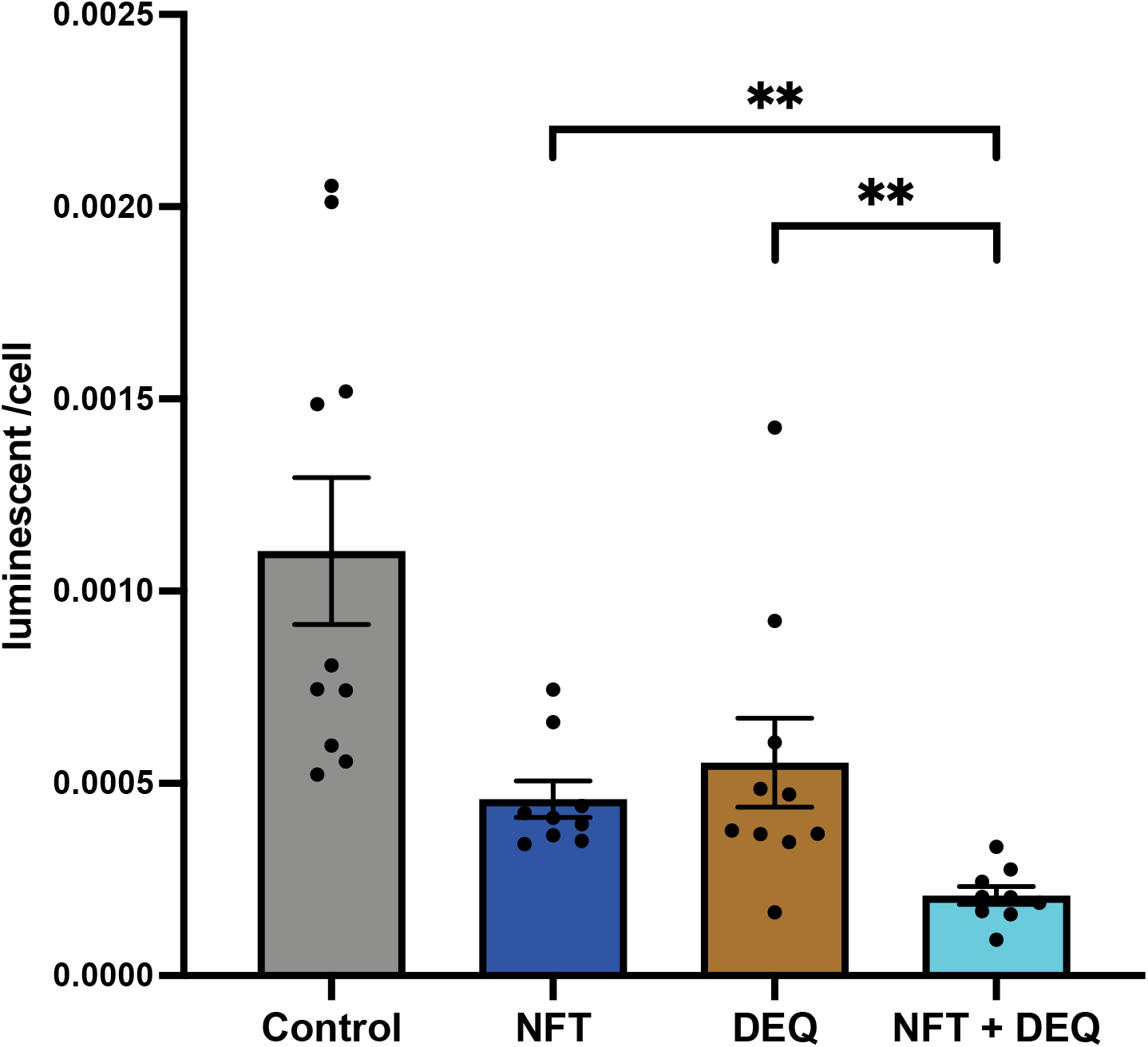
NFT + DEQ combination significantly reduces intracellular ATP levels. Intracellular ATP was quantified via luminescence in ten *E. coli* clinical isolates treated with DMSO (control), NFT, DEQ, or the NFT+DEQ combination. Experiments for each isolate were performed in triplicate, and the data represent the mean values. Data are shown as mean ± SEM. Statistical significance was determined using the Wilcoxon test. *p < 0.01.

### Dequalinium reduces persister cell formation independently of nitrofurantoin

Most antibiotics are primarily effective against actively dividing cells; in contrast, non-dividing cells can enter a dormant state that enables survival under high antibiotic concentrations [49]. These cells, known as persisters, are phenotypically tolerant rather than genetically resistant [50–52]. Given that our previous results demonstrated a significant reduction in intracellular ATP levels following treatment with NFT, DEQ, and especially their combination, we sought to determine whether ATP depletion would influence persister cell formation.

To address this, we performed a persister assay using established protocols. Briefly, we exposed bacterial cultures to antibiotic concentrations sufficient to kill most cells (and adjusted based on the drug sensitivity of the strain). After 3 or 5 hours of treatment, cells were collected, washed, and plated on LB agar to quantify surviving cells as colony-forming units per milliliter (CFU/mL).

At 3 hours post-treatment, NFT-treated cells exhibited approximately 10⁻⁴ CFU/mL persisters. In contrast, both DEQ-treated cells and cells treated with the NFT+DEQ combination showed a substantial reduction in persister levels, reaching approximately 10⁻⁶ CFU/mL. A similar trend was observed at 5 hours: NFT-treated cells yielded ∼10⁻⁵ CFU/mL persisters, whereas DEQ alone and the NFT+DEQ combination reduced persister levels further to ∼10⁻⁸ CFU/mL (Fig 4).

**Figure 4.**
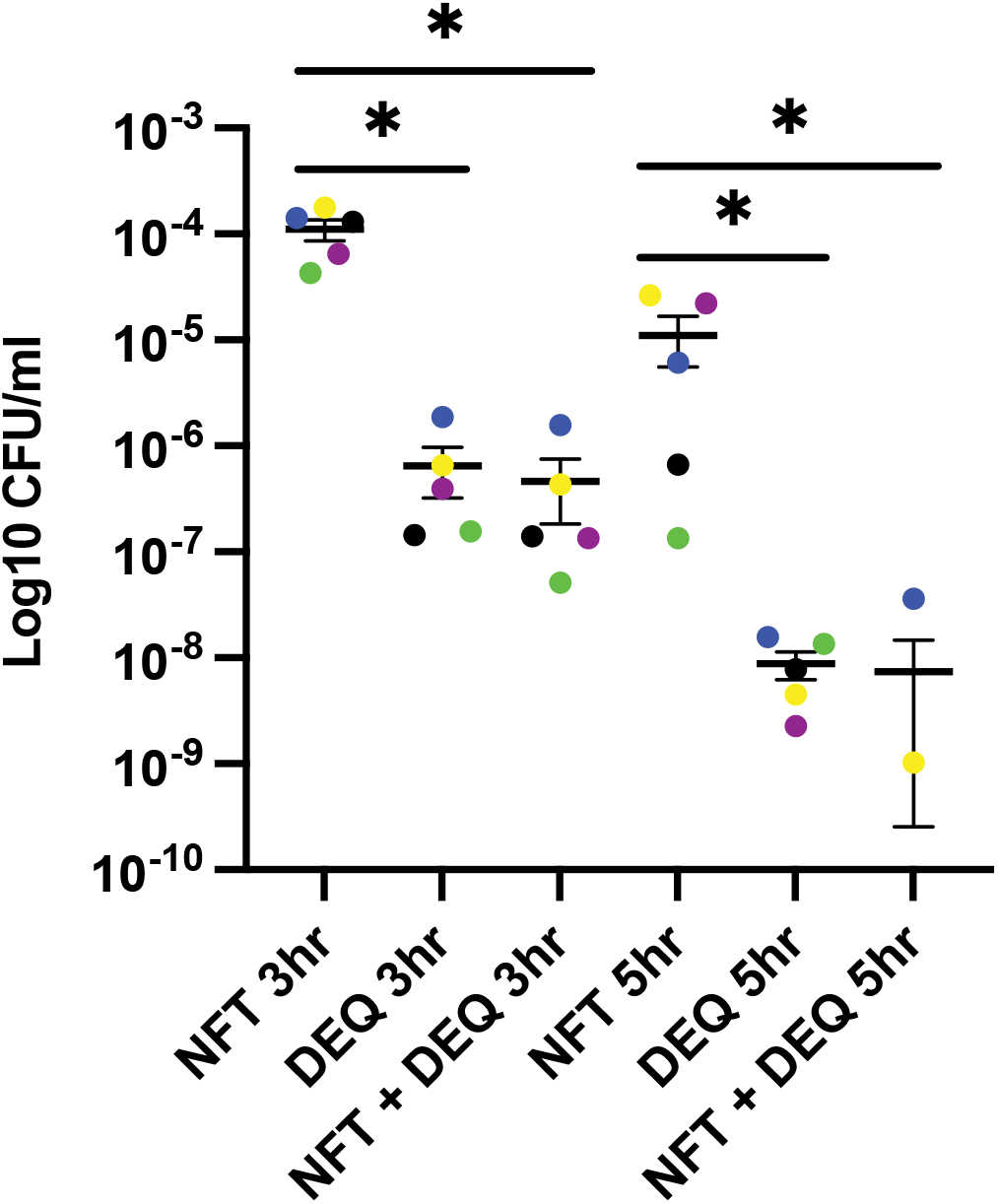
Dequalinium exposure prevents the formation of persister cells. Bacterial cultures were standardized to OD₆₀₀ = 1 before treatment with 100 µg/mL NFT or DEQ as single agents, or a combination of 50 µg/mL NFT and 50 µg/mL DEQ. Following incubation for 3 or 5 h, cells were harvested and subjected to serial dilution for colony-forming unit (CFU) quantification to assess persister cell survival. Colored dots represent different strains. Data are shown as mean ± SEM. Statistical significance was determined using the Mann-Whitney U test. *p < 0.01.

These results indicate that DEQ alone effectively reduces persister cell populations, and the addition of NFT does not further enhance this effect. Importantly, despite significant ATP depletion, the NFT+DEQ combination does not promote persister formation, suggesting that energy depletion in this context does not induce dormancy-associated tolerance. Instead, DEQ appears to counteract persister survival through mechanisms that may be independent of metabolic inhibition.

### Nitrofurantoin induces accumulation of pyruvate, citric acid and isocitrate, consistent with inhibition of TCA cycle enzymes

The specific metabolic enzyme targets of nitrofurantoin have not been well characterized. Given our observation that NFT significantly reduces intracellular ATP levels, we focused on its potential effects on central metabolic pathways, particularly the tricarboxylic acid (TCA) cycle. *E. coli* and *K. pneumoniae* are facultative anaerobes capable of growth under both aerobic and anaerobic conditions [53–55]; however, under the aerobic conditions used in this study, these organisms rely heavily on the TCA cycle for ATP production [56]. We therefore sought to identify specific enzymatic steps within this pathway that may be inhibited by NFT’s active intermediate(s).

To investigate this, we treated *E. coli* and *K. pneumoniae* cells with either DMSO (control) or NFT at the minimum inhibitory concentration (MIC) for 2 h. We subsequently washed the cells and snap-froze them in liquid nitrogen. We then partnered with the University of Utah Metabolomics Core, who extracted and analyzed the metabolites by gas chromatography–mass spectrometry (GC–MS) and performed metabolite identification using a combination of an in-house library, the NIST database, and the Fiehn metabolomics library. For *E. coli*, we analyzed both the reference strain MG1655 and a clinical isolate (BJB10); for *K. pneumoniae*, we examined two clinical isolates (KLPN1 and KLPN2).

In *E. coli* MG1655, citric acid, isocitrate, and pyruvate levels increased approximately 33-fold, 14-fold, and 5-fold, respectively, following NFT treatment. Similar trends were observed in the E. coli clinical isolate BJB10, with citric acid and isocitrate increasing 28-fold and 20-fold, respectively; however, the increase in pyruvate did not reach statistical significance (Fig. 5A). In *K. pneumoniae*, citric acid, isocitrate, and pyruvate also increased substantially, with clinical isolate KLPN1 showing 91-fold, 24-fold, and 5-fold increases, respectively, and clinical isolate KLPN2 showing 30-fold, 18-fold, and 4-fold increases, respectively. Notably, glucose-6-phosphate levels increased in *K. pneumoniae* isolates but not in E. coli (Fig. 5B).

**Figure 5.**
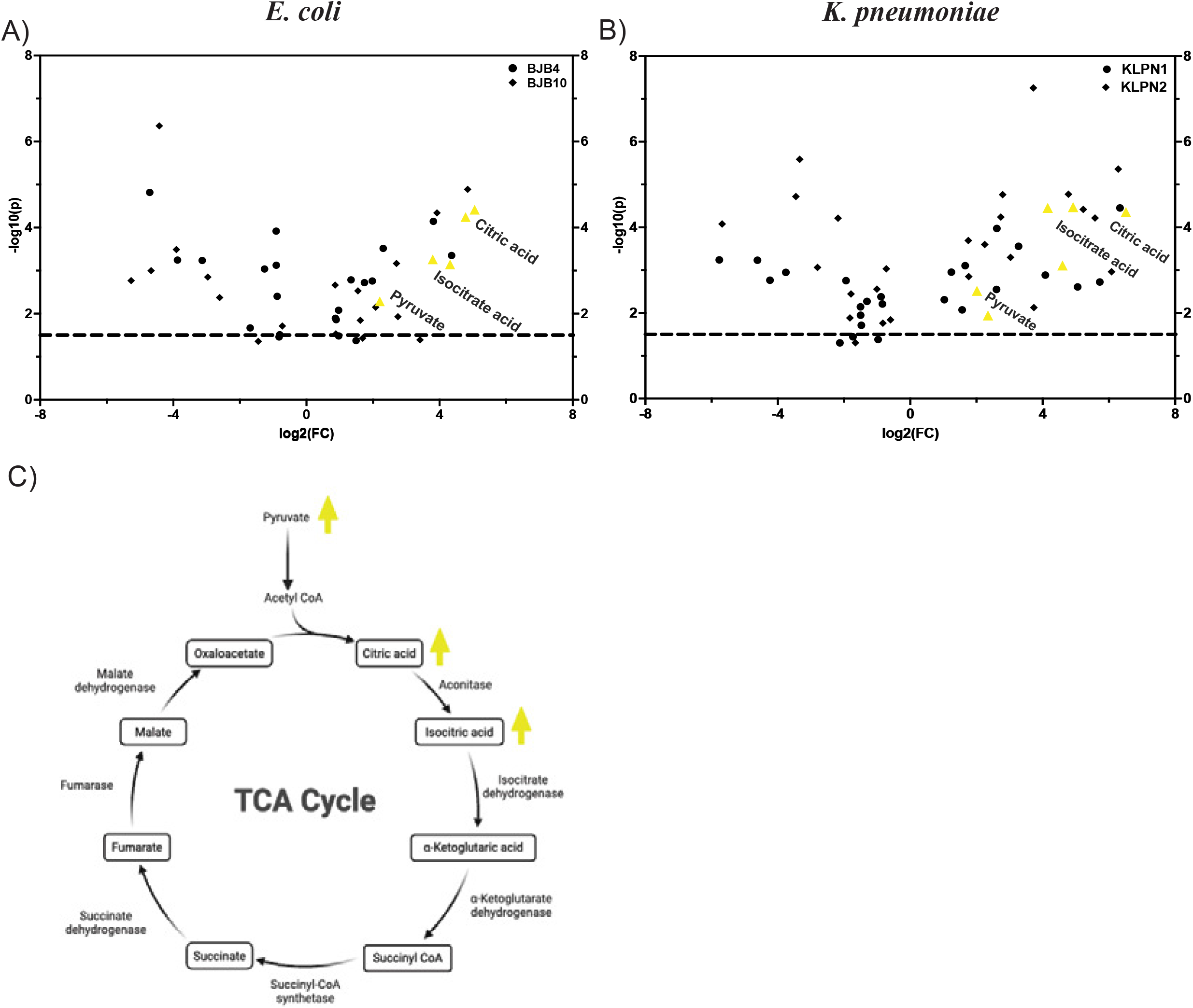
Nitrofurantoin treatment induces the accumulation of pyruvate, citric acid and isocitrate alongside the depletion of fumarate and succinate. **(A,B)** Volcano plot depicting significant shifts in metabolite abundance within *E. coli* and *K. pneumoniae*, respectively, following nitrofurantoin treatment. Circle and diamond symbols represents two different strains. Upward-pointing yellow triangles indicate metabolites that are significantly increased. Statistical analysis was performed using MetaboAnalystR. P value < 0.05 **(C)** Schematic of the TCA cycle illustrating the observed metabolic perturbations: accumulation of pyruvate, citric acid, and isocitrate (yellow upward arrows).

Within the TCA cycle, citrate is converted to isocitrate by aconitase, and isocitrate is subsequently converted to α-ketoglutarate by isocitrate dehydrogenase (57). The observed accumulation of citrate and isocitrate suggests inhibition at these steps. Furthermore, the increase in upstream glycolytic intermediates, such as pyruvate and glucose-6-phosphate, could well reflect the accumulation of precursors resulting from reduced flux into the TCA cycle (Fig. 5C).

Not all intermediates within the pathway were detected as significantly altered, which may reflect technical limitations of GC–MS detection (e.g., ionization efficiency or detection thresholds) or the rapid turnover of certain metabolites. Thus, the metabolomic data represent a snapshot of pathway perturbations rather than a complete dynamic profile. The complete list of metabolite changes is provided in Table 1 (*E. coli*) and Table 2 (*K. pneumoniae*).

**Table 1.** Significantly altered metabolites in *E. coli* following nitrofurantoin treatment used in Fig 4A.

| *E. coli* BJB4 |  |  |  |  |
| --- | --- | --- | --- | --- |
| Metabolites | FC | log2(FC) | raw.pval | -log10(p) |
| Citric acid | 33.2 | 5.1 | 0.000038 | 4.4 |
| 1-Steroylglycerol | 20.5 | 4.4 | 0.000443 | 3.4 |
| L-Threonine | 14.1 | 3.8 | 0.000071 | 4.1 |
| Isocitric acid | 13.9 | 3.8 | 0.000545 | 3.3 |
| Ribose | 4.9 | 2.3 | 0.000303 | 3.5 |
| Pyruvate | 4.6 | 2.2 | 0.005158 | 2.3 |
| Glycine | 3.9 | 2.0 | 0.001731 | 2.8 |
| 1-Palmitoylglycerol | 3.3 | 1.7 | 0.001895 | 2.7 |
| Beta-Alanine | 2.8 | 1.5 | 0.042119 | 1.4 |
| L-Lysine | 2.5 | 1.3 | 0.001633 | 2.8 |
| L-Alanine | 2.0 | 1.0 | 0.03275 | 1.5 |
| Myristic acid | 2.0 | 1.0 | 0.008304 | 2.1 |
| L-Tyrosine | 1.9 | 0.9 | 0.013791 | 1.9 |
| Capric acid | 1.8 | 0.9 | 0.012726 | 1.9 |
| L-Isoleucine | 0.6 | -0.8 | 0.030808 | 1.5 |
| Fumaric acid | 0.6 | -0.8 | 0.034636 | 1.5 |
| Threonic acid | 0.5 | -0.9 | 0.003948 | 2.4 |
| L-Glutamic acid | 0.5 | -0.9 | 0.000748 | 3.1 |
| L-Valine | 0.5 | -0.9 | 0.00012 | 3.9 |
| L-Aspartic acid | 0.4 | -1.3 | 0.000914 | 3.0 |
| L-Lactic acid | 0.3 | -1.7 | 0.021328 | 1.7 |
| 3-Phosphoglyceric acid | 0.1 | -3.1 | 0.000579 | 3.2 |
| Phosphoenolpyruvate | 0.1 | -3.9 | 0.000567 | 3.2 |
| Succinic acid | 0.0 | -4.7 | 0.000015 | 4.8 |

| *E. coli* BJB10 |  |  |  |  |
| --- | --- | --- | --- | --- |
| Metabolites | FC | log2(FC) | raw.pval | -log10(p) |
| L-Threonine | 28.8 | 4.8 | 0.000013 | 4.9 |
| Citric acid | 27.5 | 4.8 | 0.000057 | 4.2 |
| Isocitric acid | 19.9 | 4.3 | 0.00072 | 3.1 |
| 1-Steroylglycerol | 15.2 | 3.9 | 0.000046 | 4.3 |
| D-Mannose | 10.6 | 3.4 | 0.040975 | 1.4 |
| D-Fructose | 6.7 | 2.8 | 0.01182 | 1.9 |
| Beta-Alanine | 6.5 | 2.7 | 0.000686 | 3.2 |
| L-Serine | 4.2 | 2.1 | 0.007135 | 2.1 |
| Glycine | 3.2 | 1.7 | 0.037475 | 1.4 |
| Ribose | 3.1 | 1.6 | 0.014377 | 1.8 |
| 1-Palmitoylglycerol | 2.9 | 1.5 | 0.002964 | 2.5 |
| Fructose-6-phosphate | 1.8 | 0.9 | 0.029542 | 1.5 |
| L-Lysine | 1.8 | 0.9 | 0.00218 | 2.7 |
| L-Valine | 0.6 | -0.7 | 0.019531 | 1.7 |
| Nicotinamide | 0.4 | -1.4 | 0.044295 | 1.4 |
| Threonic acid | 0.2 | -2.6 | 0.004249 | 2.4 |
| 3-Phosphoglyceric acid | 0.1 | -3.0 | 0.001422 | 2.8 |
| L-Tryptophan | 0.1 | -3.9 | 0.000325 | 3.5 |
| Succinic acid | 0.0 | -4.4 | 4.35E-07 | 6.4 |
| 2-Hydroxyglutaric acid | 0.0 | -4.7 | 0.001009 | 3.0 |
| Phosphoenolpyruvate | 0.0 | -5.3 | 0.001719 | 2.8 |

**Table 2.** Significantly altered metabolites in *K. pneumoniae* following nitrofurantoin treatment used in Fig 4B.

| *K. pneumoniae* KLPN1 |  |  |  |  |
| --- | --- | --- | --- | --- |
| Metabolites | FC | log2(FC) | raw.pval | -log10(p) |
| Citric acid | 91.1 | 6.5 | 0.000044 | 4.4 |
| D-Fructose | 80.4 | 6.3 | 0.000035 | 4.5 |
| D-Glucose | 52.7 | 5.7 | 0.001894 | 2.7 |
| D-Mannose | 33.1 | 5.0 | 0.002465 | 2.6 |
| Isocitric acid | 24.1 | 4.6 | 0.000778 | 3.1 |
| Sorbitol | 16.9 | 4.1 | 0.001305 | 2.9 |
| L-Threonine | 9.6 | 3.3 | 0.000277 | 3.6 |
| Ribose | 6.1 | 2.6 | 0.000107 | 4.0 |
| 1-Steroylglycerol | 6.1 | 2.6 | 0.002823 | 2.5 |
| Pyruvate | 5.1 | 2.3 | 0.011442 | 1.9 |
| Glucose 6-phosphate | 3.1 | 1.7 | 0.000787 | 3.1 |
| 2-ketoisocaproic acid | 3.0 | 1.6 | 0.008475 | 2.1 |
| L-Serine | 2.4 | 1.2 | 0.001116 | 3.0 |
| 1-Palmitoylglycerol | 2.0 | 1.0 | 0.004909 | 2.3 |
| L-Glutamic acid | 0.6 | -0.8 | 0.006187 | 2.2 |
| L-Valine | 0.5 | -0.9 | 0.004182 | 2.4 |
| L-Lactic acid | 0.5 | -1.0 | 0.041786 | 1.4 |
| Threonic acid | 0.4 | -1.3 | 0.005386 | 2.3 |
| L-Tyrosine | 0.4 | -1.5 | 0.019467 | 1.7 |
| L-Aspartic acid | 0.4 | -1.5 | 0.011273 | 1.9 |
| Glycerol 3-phosphate | 0.4 | -1.5 | 0.007191 | 2.1 |
| Fumaric acid | 0.3 | -1.7 | 0.036004 | 1.4 |
| O-Phosphoethanolamine | 0.3 | -1.9 | 0.001764 | 2.8 |
| Nicotinamide | 0.2 | -2.1 | 0.049958 | 1.3 |
| Succinic acid | 0.1 | -3.8 | 0.001128 | 2.9 |
| 3-Phosphoglyceric acid | 0.1 | -4.2 | 0.001717 | 2.8 |
| 2-Hydroxyglutaric acid | 0.0 | -4.6 | 0.000587 | 3.2 |
| Phosphoenolpyruvate | 0.0 | -5.8 | 0.000576 | 3.2 |

| *K. pneumoniae* KLPN2 |  |  |  |  |
| --- | --- | --- | --- | --- |
| Metabolites | FC | log2(FC) | raw.pval | -log10(p) |
| L-Lysine | 78.3 | 6.3 | 0.000004 | 5.4 |
| D-Glucose | 68.0 | 6.1 | 0.001092 | 3.0 |
| D-Mannose | 48.1 | 5.6 | 0.000061 | 4.2 |
| D-Fructose | 37.6 | 5.2 | 0.000038 | 4.4 |
| Citric acid | 30.4 | 4.9 | 0.000034 | 4.5 |
| 1-Steroylglycerol | 27.6 | 4.8 | 0.000017 | 4.8 |
| Isocitric acid | 17.9 | 4.2 | 0.000035 | 4.5 |
| Sorbitol | 13.4 | 3.7 | 0.007485 | 2.1 |
| L-Threonine | 13.2 | 3.7 | 5.52E-08 | 7.3 |
| Fructose-6-phosphate | 8.2 | 3.0 | 0.000505 | 3.3 |
| Glucose 6-phosphate | 7.0 | 2.8 | 0.000017 | 4.8 |
| 1-Palmitoylglycerol | 6.7 | 2.7 | 0.000057 | 4.2 |
| Heptadecanoic acid | 4.8 | 2.3 | 0.000251 | 3.6 |
| Pyruvate | 4.1 | 2.0 | 0.003041 | 2.5 |
| Ribose | 3.4 | 1.8 | 0.001414 | 2.8 |
| L-Serine | 3.4 | 1.8 | 0.000204 | 3.7 |
| Lauric acid | 0.7 | -0.6 | 0.014473 | 1.8 |
| L-Isoleucine | 0.6 | -0.7 | 0.000931 | 3.0 |
| D-Malic acid | 0.6 | -0.8 | 0.01738 | 1.8 |
| L-Glutamic acid | 0.5 | -1.0 | 0.002771 | 2.6 |
| Inosine | 0.3 | -1.6 | 0.049809 | 1.3 |
| L-Aspartic acid | 0.3 | -1.8 | 0.003593 | 2.4 |
| Fumaric acid | 0.3 | -1.8 | 0.013052 | 1.9 |
| 2-Hydroxyglutaric acid | 0.2 | -2.2 | 0.000061 | 4.2 |
| Ethanolamine | 0.1 | -2.8 | 0.000866 | 3.1 |
| Succinic acid | 0.1 | -3.3 | 0.000003 | 5.6 |
| 3-Phosphoglyceric acid | 0.1 | -3.4 | 0.000019 | 4.7 |
| Phosphoenolpyruvate | 0.0 | -5.7 | 0.000083 | 4.1 |

Taken together, these results support a model in which NFT’s active intermediate inhibits the TCA cycle at the level of aconitase and isocitrate dehydrogenase, leading to accumulation of upstream metabolites, ultimately impairing cellular energy production.

### Isocitrate and α-ketoglutarate supplementation partially restores growth in NFT-treated cells

To test the hypothesis that NFT’s active intermediate inhibits aconitase and isocitrate dehydrogenase, we performed growth assays in the presence of exogenously supplied isocitrate, α-ketoglutarate and both isocitrate and α-ketoglutarate. We reasoned that if growth inhibition results from depletion of metabolites downstream of these enzymes, then supplementation with these intermediates should restore bacterial growth. We tested this hypothesis on *E. coli* MG1655, two clinical *E. coli* isolates, and three clinical *K. pneumoniae* isolates in a Bioscreen system with honeycomb plates. Each well contained DMSO (control) or increasing concentrations (0.5, 1, 2, or 4 mM) of either isocitrate, α-ketoglutarate or both isocitrate and α-ketoglutarate in the presence or absence of NFT. Unfortunately, because the NFT active intermediate is poorly understood and has not been synthesized or purified [38, 39], we could not add purified NFT to purified enzyme because it would not have inhibitory activity.

In the absence of NFT, isocitrate supplementation did not significantly alter growth rates in either *E. coli* or *K. pneumoniae*, as indicated by similar exponential-phase slopes across concentrations (Fig. 6A, C, and E; Fig. 7A, C, and E). Minor variations in stationary-phase OD₆₀₀ were observed but appeared to be strain-dependent rather than concentration-dependent. In the presence of NFT, isocitrate supplementation resulted in a concentration-dependent acceleration of exponential-phase growth across strains, with the most pronounced effect observed in *E. coli* isolate BJB14 (Fig. 6B) and *K. pneumoniae* isolate KLPN2 (Fig. 7B). A similar pattern was observed with α-ketoglutarate supplementation. In the absence of NFT, increasing α-ketoglutarate concentrations did not affect growth rates (Fig. 6C; Fig. 7C). However, in NFT-treated cultures, α-ketoglutarate supplementation led to a concentration-dependent acceleration of exponential-phase growth (Fig. 6D; Fig. 7D). Interestingly, in *E. coli* strain BJB14, isocitrate and α-ketoglutarate supplementation individually restored final culture density at 2 mM and further increased it at 4 mM (Fig. 6B, D), whereas in *K. pneumoniae* KLPN2, individual metabolite supplementation did not restore final culture density (Fig. 7B, D). When both isocitrate and α-ketoglutarate were added, final culture density was restored to control levels in KLPN2 (Fig. 7F) but not in BJB14 (Fig. 6F).

**Figure 6.**
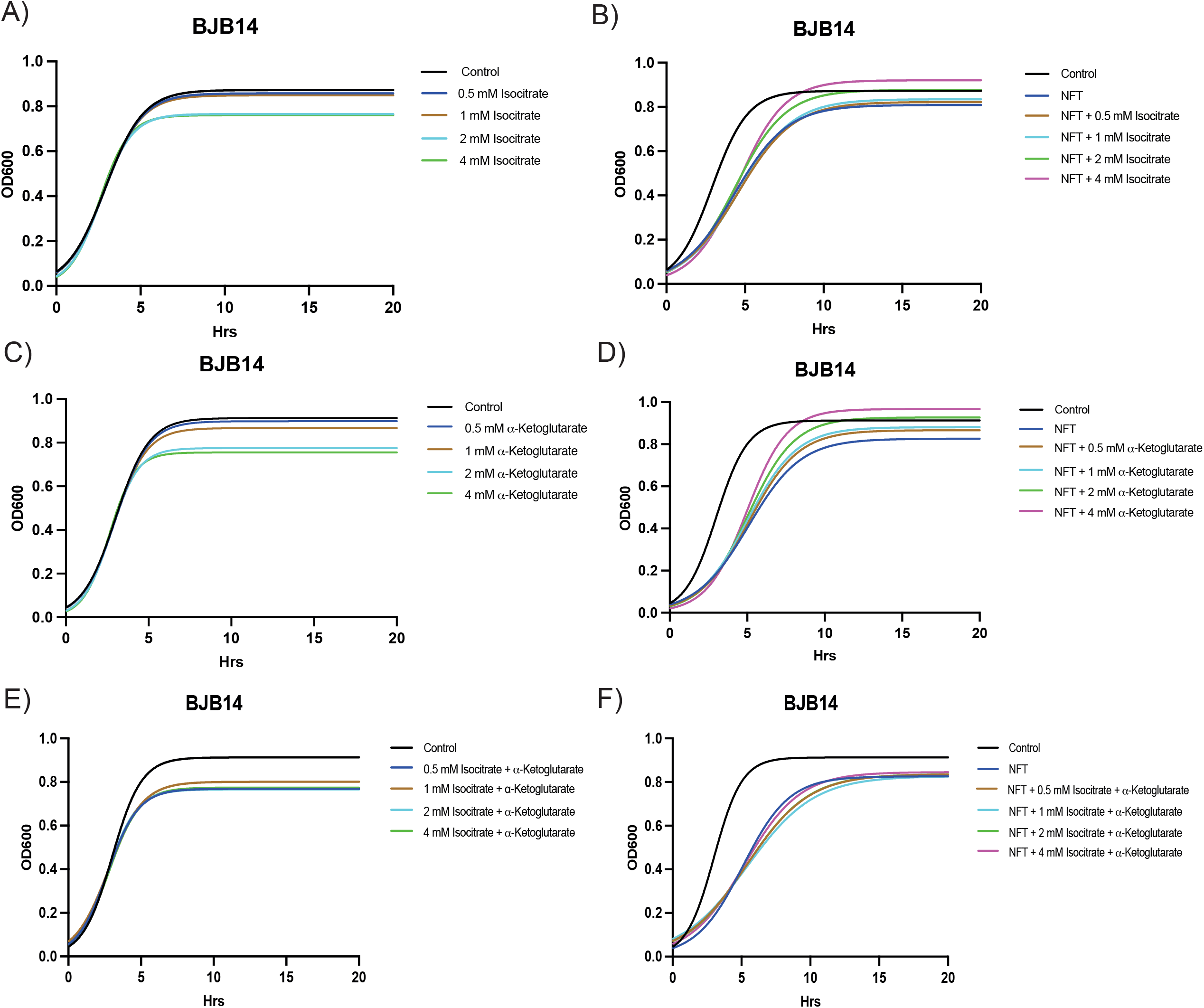
Supplementation with isocitrate, α-ketoglutarate and both metabolites partially alleviate NFT-induced growth suppression in a concentration-dependent manner. Representative growth curves are shown for *E. coli* strain BJB14, which exhibited the most prominent rescue effect. (A–B) Isocitrate supplementation, (C–D) α-ketoglutarate supplementation, and (E–F) supplementation with both metabolites. For each pair, growth curves on the left show the effect of metabolite supplementation alone (without NFT), while growth curves on the right show the effect of metabolite supplementation in the presence of NFT. Isocitrate and α-ketoglutarate supplementation alone accelerated exponential-phase growth in a concentration-dependent manner. Final culture density was restored to control levels at 2mM and increase culture density at 4mM. No rescue effect was observed upon supplementation with both metabolites combined.

**Figure 7.**
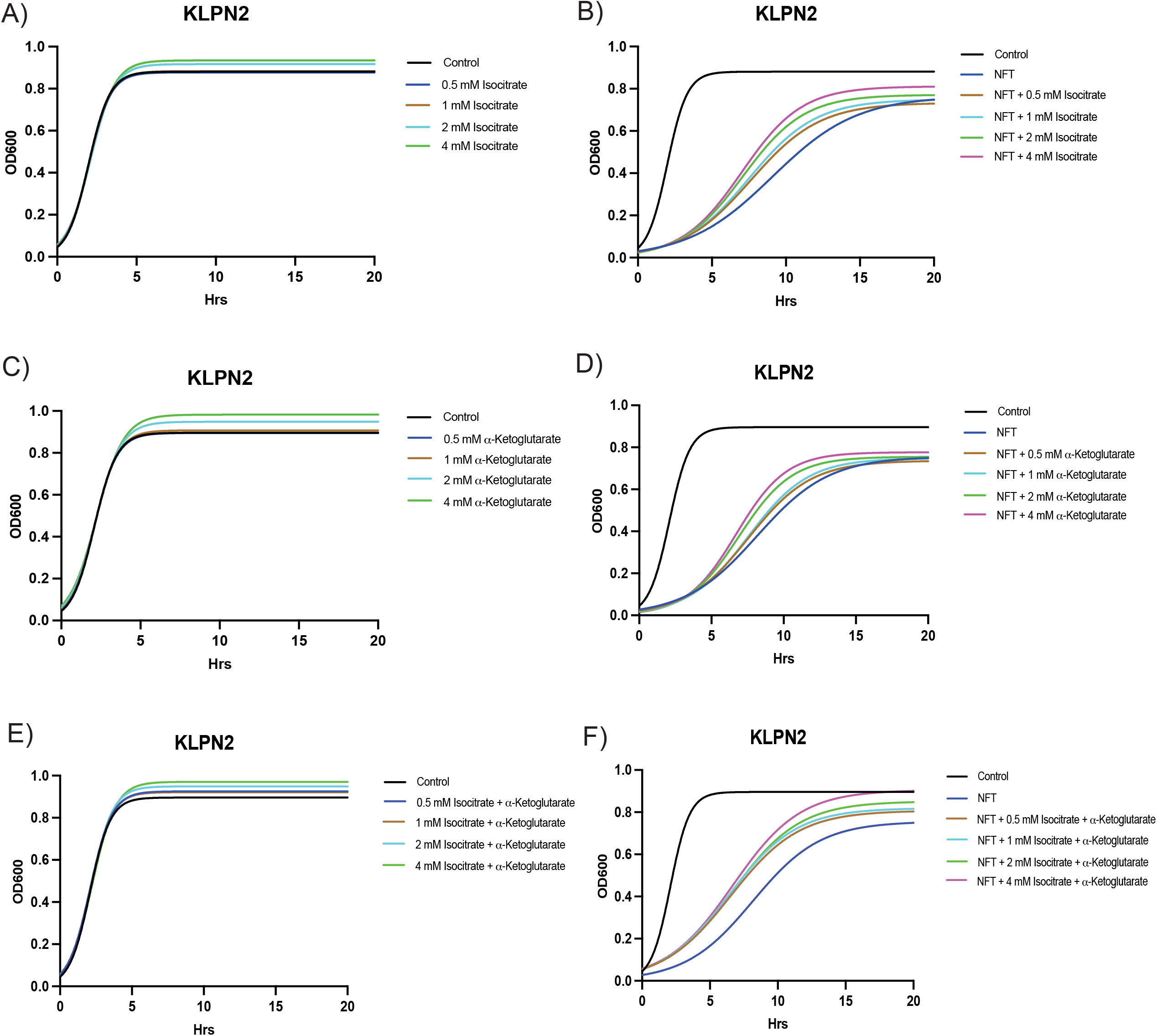
Supplementation with isocitrate, α-ketoglutarate, or both metabolites partially alleviates NFT-induced growth suppression in a concentration-dependent manner. Representative growth curves are shown for *K. pneumoniae* strain KLPN2, which exhibited the most prominent rescue effect. **(A–B)** Isocitrate supplementation, **(C–D)** α-ketoglutarate supplementation, and **(E–F)** supplementation with both metabolites. For each pair, growth curves on the left show the effect of metabolite supplementation alone (without NFT), while growth curves on the right show the effect of metabolite supplementation in the presence of NFT. As observed in *E. coli* strain BJB14, isocitrate and α-ketoglutarate supplementation alone accelerated exponential-phase growth but failed to restore final culture density to control levels. Supplementation with both metabolites also accelerated exponential-phase growth, and at 4 mM restored final culture density to control levels.

To determine whether supplementation with downstream TCA cycle metabolites would affect growth, we tested succinate supplementation and found that it inhibited exponential-phase growth and final culture density in a concentration dependent manner (Fig. S4-S5). Within the TCA cycle, intermediates serve as signaling molecules that allosterically regulate enzyme activity, and elevated metabolite levels can inhibit upstream enzymes to prevent overproduction [57, 58]. We reason that exogenous succinate may similarly trigger allosteric inhibition of upstream TCA enzymes, thereby reducing flux through the cycle and limiting energy production. Alternatively, NFT’s active intermediate may also inhibit succinate dehydrogenase; in this scenario, succinate supplementation would lead to a toxic accumulation of this metabolite. Growth curves for isocitrate-, α-ketoglutarate-, isocitrate + α-ketoglutarate-, and succinate-supplemented strains not shown in Figures 6 and 7 are included in Supplemental Figures 1–3.

Together, these supplementation experiments suggest that NFT disrupts the TCA cycle at multiple points. Supplementation with isocitrate or α-ketoglutarate appears to relieve a bottleneck upstream of these intermediates, partially restoring growth in NFT-treated cells, whereas succinate supplementation further inhibited growth, suggesting that succinate accumulation itself may be detrimental under NFT treatment. The incomplete nature of the rescue by isocitrate and α-ketoglutarate suggests that while inhibition of aconitase and isocitrate dehydrogenase likely contributes to NFT-mediated growth inhibition, additional targets or pathways are also involved.

## Discussion

Nitrofurantoin (NFT) is widely used as a first-line treatment for urinary tract infections (UTIs); however, resistance among *K. pneumoniae* approaches ∼40%, significantly limiting its clinical utility. Here, we sought to identify synergistic drug combinations capable of overcoming this resistance and to elucidate the underlying mechanisms of synergy. Through a screen of >2,500 clinically approved small molecules, we identified 21 and 15 candidates with potential synergy in combination with NFT and CPFX, respectively. We then performed broad synergy validation and confirmed that nine NFT combinations and two CPFX combinations exhibited synergy in ≥50% of strains tested. The greater number of synergistic partners identified for NFT is likely attributable to its broad and multifactorial mechanism of action, which increases the likelihood of complementary interactions, whereas CPFX primarily targets DNA gyrase and therefore presents fewer opportunities for synergistic pairing.

Among the combinations tested, the NFT–DEQ pair showed the most promise, demonstrating robust and consistent synergistic activity and offering an opportunity to repurpose the small molecule DEQ. This combination exhibited synergy in 68% of NFT-susceptible *E. coli* isolates and, notably, in 94% of NFT-resistant *K. pneumoniae* isolates, highlighting its potential to overcome clinically relevant resistance. These findings suggest that NFT–DEQ could serve as an effective therapeutic strategy against infections caused by highly resistant uropathogens.

To investigate the mechanism underlying this synergy, we focused on the hypothesis of parallel pathway inhibition. Our data indicate that NFT and DEQ cooperatively disrupt bacterial energy metabolism. Metabolomic analyses revealed that NFT treatment leads to accumulation of citrate, isocitrate and pyruvate consistent with inhibition of aconitase and isocitrate dehydrogenase in the tricarboxylic acid (TCA) cycle. This disruption likely reduces metabolic flux and limits the production of reducing equivalents required for oxidative phosphorylation. Complementary to this effect, DEQ has been reported to inhibit F1-ATPase and disrupt membrane integrity, directly impairing ATP synthesis.

Consistent with these observations, ATP quantification demonstrated that while each drug alone reduces intracellular ATP levels, the NFT+DEQ combination results in a significantly greater depletion of ATP. Functional validation through metabolite supplementation further supports this model: exogenous addition of isocitrate and α-ketoglutarate partially restored growth in NFT-treated cells, indicating that inhibition of these TCA cycle steps contributes to growth suppression. However, the incomplete rescue suggests that NFT likely affects additional cellular targets, consistent with its known pleiotropic activity.

Importantly, despite substantial ATP depletion, the NFT–DEQ combination did not increase persister cell formation. Instead, DEQ alone significantly reduced persister levels, and this effect was maintained in the combination treatment. This finding is particularly relevant, as metabolic inhibition is often associated with increased dormancy and antibiotic tolerance. Our results suggest that disruption of ATP synthesis by DEQ may counteract persister formation, thereby enhancing bacterial clearance.

Together, these data support a model in which synergy between NFT and DEQ arises from coordinated inhibition of bacterial energy metabolism (Fig. 8). Specifically, NFT disrupts ATP generation by impairing the TCA cycle, while DEQ inhibits ATP synthesis at the level of the ATP synthase. This dual targeting results in a collapse of cellular energy homeostasis, leading to bacterial killing at substantially lower drug concentrations. This mechanism may also explain the enhanced efficacy of the combination in NFT-resistant *Klebsiella spp*., as simultaneous targeting of multiple metabolic nodes can overcome resistance mechanisms that affect individual pathways.

**Figure 8.**
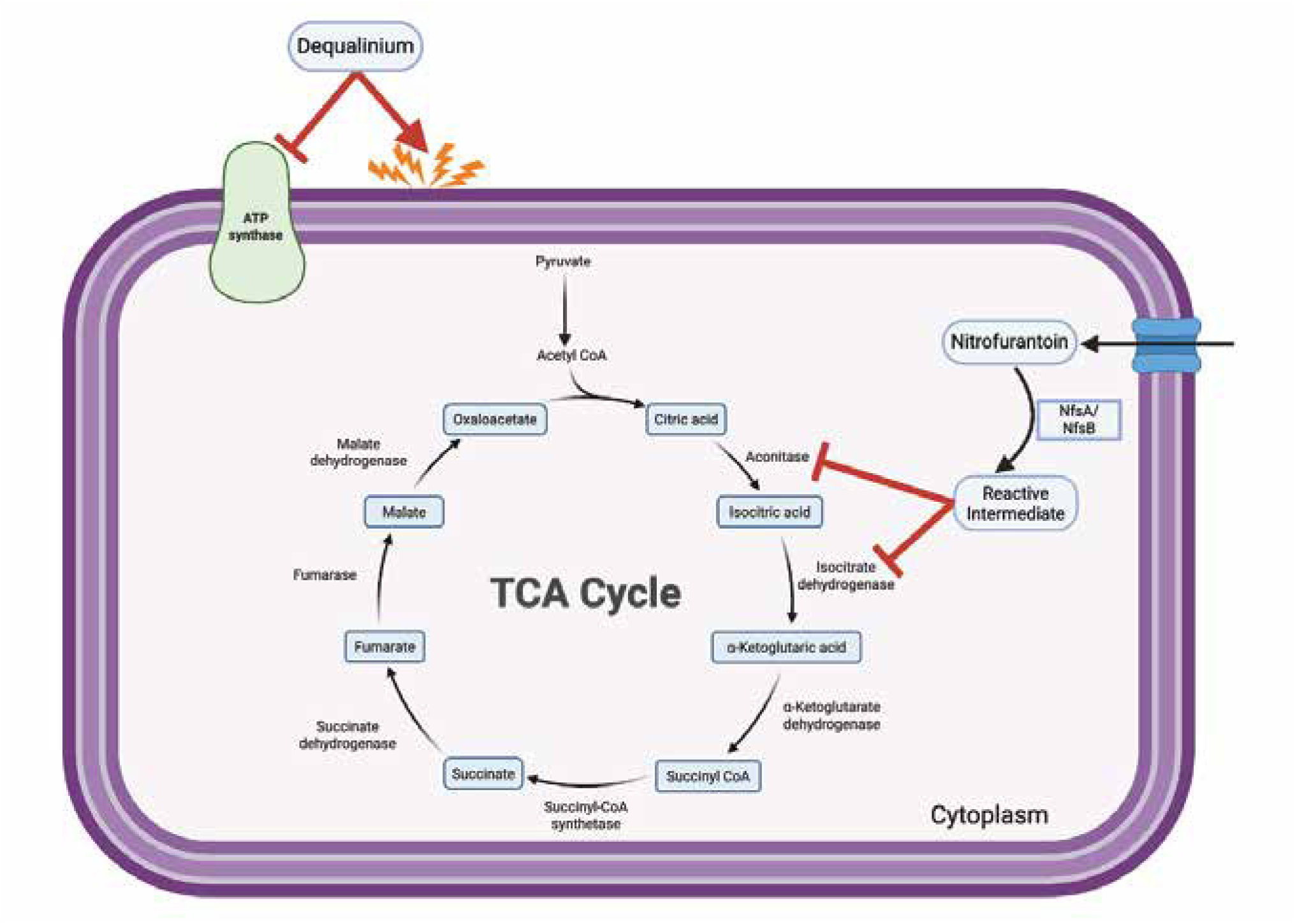
Proposed mechanism of synergy between nitrofurantoin and dequalinium. Schematic representation of dual metabolic inhibition targeting energy production. Nitrofurantoin (NFT) is taken up into the cytoplasm, where bacterial nitroreductases (NfsA and NfsB) reduce the pro-drug to a reactive intermediate. This intermediate is proposed to inhibit the TCA cycle by targeting aconitase and isocitrate dehydrogenase, leading to a metabolic bottleneck characterized by accumulation of upstream metabolites (citric acid and isocitric acid) and depletion of downstream intermediates (succinate and fumarate). Simultaneously, dequalinium (DEQ) damages the bacterial membrane and inhibits F1-ATPase (ATP synthase), directly blocking the conversion of ADP to ATP. This concurrent disruption of the TCA cycle and oxidative phosphorylation results in depletion of intracellular ATP levels.

In summary, our study identifies the NFT–DEQ combination as a promising therapeutic strategy for treating drug-resistant UTIs and provides mechanistic insight into its synergistic activity. More broadly, these findings highlight the potential of targeting bacterial energy metabolism through combination therapy as an effective approach to combat antibiotic resistance.

## Materials and methods

### Bacterial strains

The initial drug screen was performed using the *E. coli* reference strain MG1655. Synergy validation, checkerboard assays, metabolomic analyses, and subsequent experiments were conducted using *E. coli* MG1655 and clinical isolates designated BJB#, where # indicates the isolate number. *K. pneumoniae* strains used in this study were exclusively clinical isolates, designated KLPN#, where # indicates the isolate number. All clinical isolates were obtained from patients with positive urine cultures and were identified and archived by the microbiology laboratory at Primary Children’s Hospital (Salt Lake City, UT, USA).

### Minimum inhibitory concentration (MIC) assays

MIC assays were performed in M9 minimal medium composed of 10.5 g/L M9 broth (Amresco), 0.2% casamino acids, 0.4% glucose, 0.1 mM CaCl₂, 1 mM MgSO₄, 0.25% nicotinic acid, and 0.33% thiamine in H₂O, unless otherwise specified. Overnight cultures were grown at 37 °C in a roller drum and subsequently diluted to an OD₆₀₀ of 0.002.

Each well was inoculated with approximately 1 × 10³ cells (2 µL of diluted culture into 200 µL of medium per well). Plates were incubated at 37 °C for 24 h. Small molecules were dissolved in either DMSO or Milli-Q H₂O and subjected to twofold serial dilution.

Optical density at 600 nm (OD₆₀₀) was measured using a SpectraMax iD5 plate reader (Molecular Devices, San José, CA, USA) immediately after inoculation and following 24 h of incubation. MIC values were defined as the lowest concentration of compound resulting in ≥90% inhibition of growth relative to untreated controls.

### High-Throughput Drug Screen

The drug screen was conducted using three 96-well plates, with each well containing a single small molecule from the Spectrum Library Collection in combination with the antibiotics of interest, nitrofurantoin (NFT) or ciprofloxacin (CPFX). Compounds from the library were diluted from a 1 mM stock to a working concentration of 20 µM. Sub-inhibitory concentrations of each antibiotic, corresponding to 2-fold and 4-fold below the minimum inhibitory concentration (MIC), were prepared separately.

Plate 1 contained 50 µL of the 20 µM small-molecule solution and 50 µL of antibiotic at 2-fold below MIC. Plate 2 contained 50 µL of the 20 µM small-molecule solution and 50 µL of antibiotic at 4-fold below MIC. Plate 3 served as a control and contained 50 µL of the 20 µM small-molecule solution without antibiotic. Following inoculation, the final concentration of each small molecule was 0.4 µM in all wells.

Overnight bacterial cultures were grown at 37 °C and diluted to an OD₆₀₀ of 0.002. Each well was inoculated with 2 µL of the diluted culture. Plates were incubated at 37 °C for 24 h, after which optical density at 600 nm (OD₆₀₀) was measured using a SpectraMax iD5 plate reader (Molecular Devices, San José, CA, USA). Growth inhibition of ≥90% relative to control wells was used to identify potential synergistic interactions, consistent with the criteria used for MIC determination.

### Checkerboard Assay and Fractional Inhibitory Concentration Index (FICI) Calculation

Overnight cultures were grown at 37 °C and diluted to an OD₆₀₀ of 0.002 prior to use. Checkerboard assays were performed in 96-well plates as previously described [20,21]. Briefly, Drug A was serially diluted across columns 1–11, and Drug B was serially diluted across rows A–G. Column 12 and row H served as single-drug controls. Each well was inoculated with 2 µL of the diluted bacterial culture. Plates were incubated at 37 °C for 24 h, and optical density at 600 nm (OD₆₀₀) was measured using a SpectraMax iD5 plate reader (Molecular Devices, San José, CA, USA) both immediately after inoculation and after incubation.

Drug interactions were evaluated by calculating the fractional inhibitory concentration index (FICI) using the following formula:

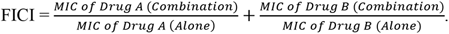

A FICI value ≤ 0.5 was defined as synergistic. All checkerboard assays were performed in triplicate for each strain–drug combination. When results were consistent (i.e., all replicates classified as synergistic or non-synergistic), the mean FICI value was calculated and used for data presentation.

### ATP Quantification Assay

Intracellular ATP levels were measured using the BacTiter-Glo™ Microbial Cell Viability Assay (Promega Corporation, Madison, WI, USA) according to the manufacturer’s instructions. Overnight cultures were grown at 37 °C and subsequently diluted into fresh M9 medium, followed by incubation for ∼2 h to restore cells to logarithmic-phase growth.

Cells were then treated with the minimum inhibitory concentrations (MICs) of nitrofurantoin (NFT) or dequalinium (DEQ). For the combination treatment, a concentration corresponding to 4-fold below the MIC of each drug was used to represent synergistic conditions. DMSO-treated cells served as the negative control. After 1 h of treatment, 100 µL of culture was transferred into three replicate wells of a 96-well plate.

An equal volume of BacTiter-Glo™ reagent was added to each well, and plates were mixed for 15 s using the orbital shaking function of a SpectraMax iD5 plate reader (Molecular Devices, San José, CA, USA). Plates were then incubated at room temperature for 5 min before luminescence was measured. Luminescence values were averaged across replicates and normalized to OD₆₀₀ to obtain ATP levels on a per-cell basis.

### Metabolomics

Overnight cultures were grown at 37 °C. The following day, 500 µL of culture was inoculated into 50 mL of fresh M9 medium. Cultures were incubated at 37 °C with shaking for approximately 2 hours to allow recovery to mid-log phase. Cells were then exposed for 1 hour at 37 °C with shaking to either DMSO (negative control) or the MIC of nitrofurantoin. Following treatment, cells were harvested by centrifugation, washed with PBS, and flash-frozen in liquid nitrogen. Samples were subsequently submitted to the University of Utah Metabolomics Core Facility for processing and analysis. Statistical analysis was performed using MetaboAnalystR [59].

### Metabolites Supplementation Growth Curve

Growth curves were generated using a Bioscreen G Pro instrument (BIOSCREEN, Turku, Finland) with manufacturer-specific 100-well honeycomb plates. Cultures were prepared in 1.7 mL microcentrifuge tubes, and 200 µL of each condition was dispensed into individual wells.

Plates were incubated at 37 °C with continuous shaking for 24 h, and optical density at 600 nm (OD₆₀₀) was recorded at 30 min intervals. Each condition was assayed in quadruplicate, and mean values were used for subsequent analysis.

### Persister Assay and CFU Enumeration

Overnight cultures were grown at 37 °C and normalized to an OD₆₀₀ of 1.0. Subsequently, 50 µL of normalized culture was inoculated into 5 mL of fresh M9 medium and incubated at 37 °C with shaking until mid-log phase (∼2 h). At mid-log phase, cultures were treated with nitrofurantoin (NFT), dequalinium (DEQ), or a combination of both. Final concentrations were 100 µg/mL for single-drug treatments and 50 µg/mL of each compound for the combination treatment. Cultures were then incubated at 37 °C with shaking. At 3 and 5 h post-treatment, 1 mL aliquots were collected and pelleted by centrifugation. Cells were washed with cold phosphate-buffered saline (PBS) to remove residual antibiotics and halt further growth. Samples were serially diluted (10⁰–10⁻³) and plated on LB agar for colony enumeration. Plates were incubated at 37 °C for 24 h, after which colony-forming units (CFUs) were quantified.

## Acknowledgments

This work was supported by Congressionally Directed Medical Research Program grant W81XWH-22-1-0800 (SC210103) from the Department of Defense. Special thanks to the Metabolomics Core at The University of Utah for performing the metabolomics experiments and analysis.

---

**Supplemental Figure 1. Growth curves with isocitrate supplementation.** *E. coli* strains BJB4 and BJB10 (A–D) and *K. pneumoniae* strains KLPN1 and KLPN12 (E–H). Growth curves on the left show isocitrate supplementation alone, without NFT (A, C, E, G), while growth curves on the right show isocitrate supplementation in the presence of NFT (B, D, F, H).

**Supplemental Figure 2. Growth curves with α-ketoglutarate supplementation.** *E. coli* strains BJB4 and BJB10 (A–D) and *K. pneumoniae* strains KLPN1 and KLPN12 (E–H). Growth curves on the left show α-ketoglutarate supplementation alone, without NFT (A, C, E, G), while growth curves on the right show α-ketoglutarate supplementation in the presence of NFT (B, D, F, H).

**Supplemental Figure 3. Growth curves with isocitrate and α-ketoglutarate supplementation.** *E. coli* strains BJB4 and BJB10 (A–D) and *K. pneumoniae* strains KLPN1 and KLPN12 (E–H). Growth curves on the left show isocitrate and α-ketoglutarate supplementation alone, without NFT (A, C, E, G), while growth curves on the right show isocitrate and α-ketoglutarate supplementation in the presence of NFT (B, D, F, H).

**Supplemental Figure 4. Growth curves with succinate supplementation.** *E. coli* strains BJB4, BJB10, and BJB14 (A–F). Growth curves on the left show succinate supplementation alone, without NFT (A, C, E), while growth curves on the right show succinate supplementation in the presence of NFT (B, D, F).

**Supplemental Figure 5. Growth curves with succinate supplementation.** *K. pneumoniae* strains KLPN, KLPN2 and KLPN12 (A–F). Growth curves on the left show succinate supplementation alone, without NFT (A, C, E), while growth curves on the right show succinate supplementation in the presence of NFT (B, D, F).

